# Sex specific systemic effects of *sev-Gal4* driven activated Ras expression mediated through hnRNPs in *Drosophila*

**DOI:** 10.64898/2026.02.27.708457

**Authors:** Vanshika Kaushik, Subhash C. Lakhotia

## Abstract

Following our laboratory’s earlier observations on systemic damage inflicted by *sev-Gal4* driven activated Ras (*sev*>*Ras*^*V12*^) over-expression in *Drosophila* larval eye discs, we now show that *sev*>*Ras*^*V12*^ expressing males suffer enhanced eye roughening and pupal death than female sibs because the former have significantly greater Ras levels in ommatidial cells than in female counterpart. In normally developing ommatidial cells, TBPH/TDP-43 was more abundant in cytoplasm in male than in female eye discs. The *sev*>*Ras*^*V12*^ expression reduced nuclear TBPH in female eye discs but caused no apparent change in males. Caz/Fus, an interacting partner of TBPH, was significantly downregulated in *sev*>*Ras*^*V12*^ eye discs, more so in males. Significant reduction in the microtubule binding protein Futsch in eye discs of *sev*>*Ras*^*V12*^ larvae of either sexes but female-specific elevation of Fas2 appears to be due to the above normal developmental differences in TBPH and Caz in female and male ommatidial cells and because Sxl, the master regulator of sex-determination, is present only in females. In view of known auto-regulatory loop between Fas2 and Ras, we suggest that elevated levels of Fas2 cause levels of Ras to be much less elevated in *sev*>*Ras*^*V12*^ female eye discs than in male sibs. This results in greater local and systemic damage in males. These findings have general and clinical relevance since perturbed Ras signaling is a major factor in several diseases, including cancer.

## 1. Introduction

An earlier study from our laboratory [1] found that over-expression of activated Ras^V12^ in *sevenless* expressing cells (*sev*>*Ras*^*V12*^) of developing eye discs in *Drosophila* larvae not only causes eye-roughening but also has a systemic effect culminating in extensive pupal death [2]. These studies also showed that mis-expression of the lncRNAs encoding *hsr*ω gene in *sev*>*Ras*^*V12*^ background exaggerates the eye-roughening and pupal lethality phenotypes. While studying interactions between *hsr*ω lncRNAs and Ras^V12^ over-expression, we noted that the rough eye and pupal lethality phenotypes of *sev*>*Ras*^*V12*^ were significantly more pronounced in male than in female sibs and that the exaggeration of these phenotypes by mis-expression of *hsr*ω transcripts was also sex-biased with males being more severely affected.

Sex-biased effects of many mutant and pathogenic conditions have been widely noted in diverse organisms [3-9]. The sex-biased differential responses, which span from general behaviour to metabolism, nutrient storage and transport, response of neural tissues and disease propensity, are mostly be associated with the sex-determination pathways indicating that the organism’s gender dictates how it behaves in and responds to altered external and internal environments.

A transcriptomic analysis in the earlier study [1] suggested that the heterogenous RNA-binding proteins (hnRNPs) have important role in mediating the interaction between Ras signaling and *hsr*ω. Diverse hnRNPs are widely involved in nucleic acid metabolism including splicing, RNA transport and stabilization, DNA repair, and regulation of translation and transcription [10-14]. In the present study we examined interactions of three of the key hnRNPs, viz., TBPH/TDP-43, Caz /dFus and Sxl, in *sev*>*Ras*^*V12*^ background. Our results suggest that the greater damage in *sev*>*Ras*^*V12*^ expressing eye discs of male larvae follows differential interactions between these and other hnRNPs in the two sexes.

## 2. Material and methods

### 2.1. Fly stocks

All the stocks and crosses were maintained on standard agar cornmeal medium at 24±1 °C. The following fly stocks were used; Bloomington Drosophila Stock Ctr (BDSC) numbers are given, where applicable:

*w*^*1118*^; *sev-Gal4;+*/*+* (BDSC no. 5493)

*w*^*1118*^; *UAS-eGFP;+*/*+* (BDSC no. 5431)

*w*^*1118*^; *+*/*+; UAS-Ras*^*V12*^ (BDSC no. 4847)

*w*^*1118*^; *UAS-TBPH-RNAi/CyO; +*/*+* (BDSC no. 39014, [15]; from Girish Ratnaparakhi, Pune

*w*^*1118*^; *UAS-TBPHGFP; +*/*+* expresses GFP-tagged TBPH under UAS control; from Daniella Zarnescu, University of Arizona [16]

*w*^*1118*^; *+*/*+; UAS-cazRNAi* (BDSC no. 34839); from Girish Ratnaparakhi, Pune

*w*^*1118*^; *+*/*+; UAS-FUS*^*WT*^ *-* expresses HA-tagged wild type human FUS, a functional homolog of *Drosophila* Caz [17], under UAS control [15]; from Uday Bhan Pandey, Louisiana State University

*w*^*1118*^; *+*/*+; UAS-hsr*ω*-RNAi*^*3*^, targets the 280bp tandem repeats in the *hsr*ω*RB hsr*ω*RG*, and *hsr*ω*RF* lncRNAs [18]

*w*^*1118*^; *+*/*+; 3x Atg8a-mcherry*, expresses mcherry-tagged Atg8a under endogenous promoter [19]; from Bhupendra Shravage, Agarkar Research Institute, Pune.

### 2.2. Lethality Assay

Crosses between flies of the desired genotypes were set up at 24±1 °C and synchronized progenies were collected. Data for total numbers of embryos, hatched 1^st^ instar larvae, larvae that pupated, and numbers of pupae that eclosed were collected from at least two biological replicates for each genotype.

### 2.3. Photomicrography of adult eyes

For examining the external eye morphology of flies of the desired genotypes, eclosed flies were etherized and their eyes were photographed using a Zeiss Axiocam HRc Camera mounted on Nikon SMZ800N stereobinocular microscope.

### 2.4. Immunostaining

Eye discs of 120 hr male and female larvae of desired genotypes were dissected in 1xPBS (Phosphate Buffer Saline, 1.37 M NaCl, 27 mM KCl,100 mM Na2HPO4, and 20mM KH2PO4, pH7.4) followed by fixation in 4% freshly prepared Paraformaldehyde in 1X PBS for 20 min followed by immunostaining as described earlier [20]. Briefly, fixed tissues were washed in 0.1% PBST (1X PBS with 0.1% Triton-X) 3 times for 15 min each followed by incubation in blocking solution (0.1% Triton X-100, 0.1% BSA, 10% FCS, 0.1% deoxycholate, and 0.02%Thiomersal) for 2 hr at room temperature, incubation in the desired primary antibody for 2 hr at room temperature, washing with 0.1% PBST 3 times, blocking, and finally incubation in specific secondary antibody followed by 3 times washing with 0.1% PBST, DAPI staining (4′,6-Diamidino-2-Phenylindole, Dihydrochloride, Thermo Fisher Scientific, Cat# D1306, 1 μg/ml) for 30 min, 0.1% PBST washing and finally, mounting in DABCO (1,4-Diazabicyclooctane, Sigma, Cat# D27802). Following primary antibodies were used: rabbit monoclonal anti-Ras (27H5, Cell signaling, 1:100), rabbit polyclonal anti-TBPH (10782-2-AP, Proteintech, 1:100), mouse monoclonal anti Caz (3F4,1:100, provided by Moushami Mallik, Germany [21], mouse monoclonal anti-Sxl (M114-s, Developmental Studies Hybridoma Bank, 1:20), mouse anti-Futsch/22c10-s (AB_528403, Research Resource Identifier, 1:20), and mouse monoclonal anti-FasII (1D4-s, Developmental Studies Hybridoma Bank, 1:20). Following secondary antibodies from Invitrogen were used: Goat anti-mouse Alexa-Fluor 647, Goat anti-mouse Alexa-Fluor 546, Donkey anti-Rabbit Alexa-Fluor 546, Goat anti-Rabbit Alexa-Fluor 633 (Invitrogen); goat anti-mouse Alexa-Fluor 488 was from Jakson Immuno Research.

### 2.5. Microscopy and image processing

The immunostained preparations were examined using Zeiss LSM-510 Meta at Department of Zoology, BHU, Zeiss LSM-900 at Dr. Bama Charan Mondal’s Lab, Department of Zoology, BHU, Zeiss LSM-510 at Interdisciplinary School of Life Sciences (ISLS), BHU, or Leica SP8 STED facility at Central Discovery Centre (CDC), BHU. Fiji/ImageJ software (NIH, USA) and Zen Blue (Zeiss) softwares were used to analyse and quantify images. Images were arranged using Adobe photoshop 2020.

Fluorescence intensity in confocal images was quantified from Maximum intensity projection images of the proximal posterior regions in three consecutive optical sections of eye discs showing the R7 photoreceptors, identified by the *sev*>*eGFP* fluorescence, were obtained. For total fluorescence intensity quantification, a circular region of interest (ROI) was drawn and the total intensities of the ROI were measured. For quantifying TDP-43 intensity in R7 nuclei, 3 R7 nuclei were manually marked using freehand selection tool of ImageJ in each disc to obtain the average TDP-43 intensity in R7 nuclei in the given eye disc. For TDP-43 intensity profile, Zen Blue software was used to draw a line across R7 to R3/R4 cell nuclei in single optical section to plot the intensity profile graph. Fluorescence intensity graphs were plotted and statistical test was performed using Graph-Pad Prism 8.4.2 software wherein a 2-tailed unpaired *t*-test was performed to evaluate the statistical significance between mean of 2 groups. In each graph significance is indicated by * for p≤0.05, ** for p≤0.01, *** for p≤0.001, and **** for p≤0.0001 and ‘ns’ for non-significant difference.

## 3. Results

### 3.1. *Sev-Gal4* driven expression of Ras^V12^ results in sex biased eye roughening and pupal lethality and higher levels of Ras in male eye imaginal discs

Expression of the constitutively active Ras^V12^ under *sev-Gal4* driver (*sev* > *Ras*^*V12*^) in eye discs causes rough eye and late pupal lethality [1]. Our present study revealed that the male and female *sev* >*Ras*^*V12*^ progenies were differentially affected (Figs. 1a-e), with males exhibiting greater eye roughening and necrotic patches (Fig.1c, d). Further, more *sev* >*Ras*^*V12*^ male than female pupae died since out of the approximately 40% of *sev*>*Ras*^*V12*^ pupae that eclosed, only about 10% were male (Fig.1e). In agreement with earlier report [1] most *sev*>*Ras*^*V12*^ *hsr*ω^*RNAi*^ progenies, irrespective of gender, were seen to die as early pupae with none eclosing (Fig. 1e).

**Fig. 1.**
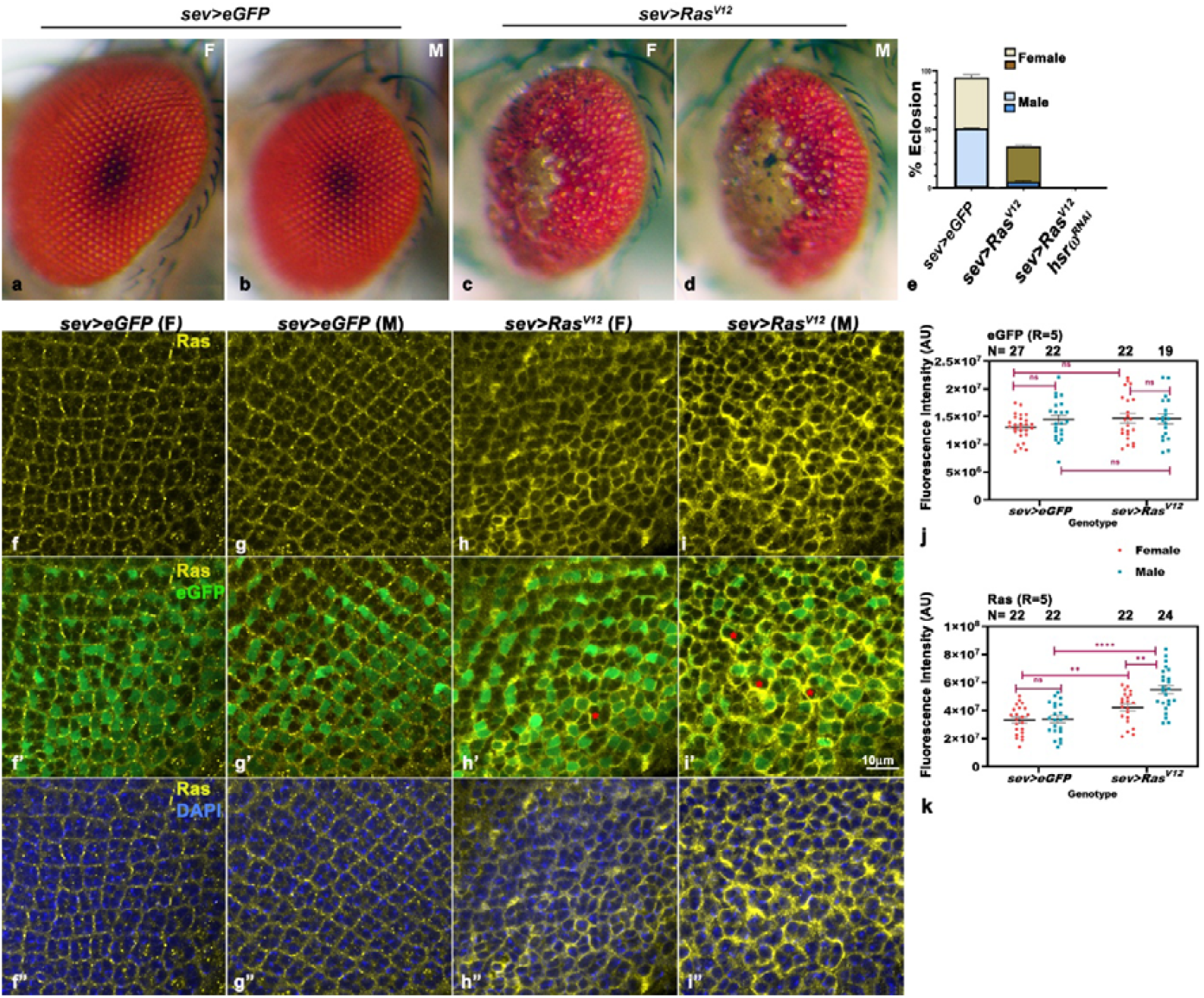
Sex-biased effects of *sev*>*Ras*^*V12*^ expression on eye-roughening, pupal death and Ras levels in larval eye discs. **a-d**. Photomicrographs of eyes of female (**a, c**) and male adults (**b, d**) of the indicated genotypes (top). **e**. Bar graphs showing mean (± S.E.; data from 3 independent replicates) percent eclosion (Y-axis) of *sev*>*eGFP* (N=720), *sev*>*Ras*^*V12*^ (N=825) and *sev*>*Ras*^*V12*^ *hsr*ω^*RNAi*^ (N=607) pupae. **f-i”** Confocal optical sections of posterior region of eye discs of 120 hr control *sev*>*eGFP* (**f-g”**) and *sev*>*Ras*^*V12*^ (**h-i”**) female (**F**) and male (**M**) larvae showing localization of Ras (yellow), eGFP (green) and nuclei (DAPI, blue); red asterisks (**\***) in **h’** and **I’** mark some of the non-GFP expressing cells which have high levels of Ras. Scale bar in **i** = 10µm, and applies to **f-i”** images. **j-k**. Dot plots showing mean (± S.E.) fluorescence intensity (Y-axis in arbitrary units (AU)) of eGFP (**j**) and Ras (**k**), respectively, in different genotypes (X-axis); **R** in **j** and **k** refers to numbers of biological replicates; the total numbers (**N**) of discs examined in each genotype are indicated by the numbers above each dot-plot column.

Immunostaining of *sev*>*eGFP* and *sev*>*Ras*^*V12*^ eye discs revealed that while the Ras levels in *sev*>*eGFP* male and female eye discs were comparable (Fig. 1f-g”), the *sev-Gal4* driven overexpression of the *Ras*^*V12*^ transgene resulted in significantly greater Ras levels in *sev*>*Ras*^*V12*^ male, than in female, eye discs (Fig. 1h-i”). To check if the *sev-Gal4* driver was more active in males, which may cause an enhanced expression of the *Ras*^*V12*^ transgene, we quantified eGFP and Ras fluorescence signals in a 27.343 µm radius circular ROI in projection images of the proximal posterior region of 3 optical sections in which the R7 was distinctly identifiable. While the eGFP signal in Ras^V12^ over-expressing female and male eye discs (Fig. 1j) remained similar to that in the *sev-eGFP* controls, the Ras staining in same regions in Ras^V12^ over-expressing eye discs showed a significantly greater increase in male than in female eye discs (Fig. 1k). It is notable that in agreement with the earlier report [1] of non-cell autonomous spread of Ras expression, we found that the increase in Ras levels extended also to the GFP-ve cells that do not express the *sev-Gal4* driver, more so in male discs (Fig. h-i’’).

### 3.2. Co-alteration of TBPH or Caz levels enhance *sev*>*Ras*^*V12*^ pathogenicity but marginally reduce the exaggerated pupal lethality of *sev*>*Ras*^*V12*^ *hsr*ω^*RNAi*^ pupae

In agreement with an earlier study from our laboratory [1], we also observed complete pupal lethality when *hsr*ω transcripts were down-regulated in Ras^V12^ over-expression background (Fig. 1). To understand the gender-biased phenotypes following *sev-Gal4* driven *Ras*^*V12*^ expression and their exaggeration by down-regulation of *hsr*ω transcripts, we examined effects of altered levels of TBPH/TDP-43 and Caz/dFus in these genetic backgrounds. These two key hnRNPs are known to interact with Ras signaling as well as with *hsr*ω transcripts. Activated MAPK, a downstream effector of Ras signaling, phosphorylates TDP-43 at known phosphorylation sites of TDP-43 i.e.,Thr-153/Tyr-155 [22] while hyperactivated MAPK hyperphosphorylates TDP-43 at these two sites affecting its RNA binding affinity and splicing activities [23, 24]. Caz negatively regulates EGFR signaling [25]. Both the hnRNPs physically interact with the *hsr*ω lncRNAs in omega speckles (Piccolo et al., 2014), which store and regulate dynamics of different hnRNPs and some other RNA binding proteins [26, 27]. TDP-43 and Caz also physically interact with each other and have common downstream targets [28-31].

Down-regulation of TDP-43 in *sev*>*Ras*^*V12*^ background substantially enhanced lethality at late pupal stage with very few adult females, but no males, emerging (Fig. 2a). The rare emerging *sev*>*Ras*^*V12*^ *TDP-43*^*RNAi*^ females had very severely damaged eyes (Fig. 2b). Total pupal lethality, adult emergence and eye damage in *sev*>*Ras*^*V12*^ *TDP-GFP* progeny was comparable to that in *sev*>*Ras*^*V12*^, except that ∼30% *sev*>*Ras*^*V12*^ *TDP-GFP* pupae died as early pupa (Fig. 2a, d, e). Interestingly, compared to 100% death of *sev*>*Ras*^*V12*^ *hsr*ω^*RNAi*^ as early pupae, up-or down-regulation of TBPH in *sev*>*Ras*^*V12*^ *hsr*ω^*RNAi*^ background permitted some pupae to develop till mid and late stages, although none of them emerged as fly (Fig. 2a).

**Fig. 2.**
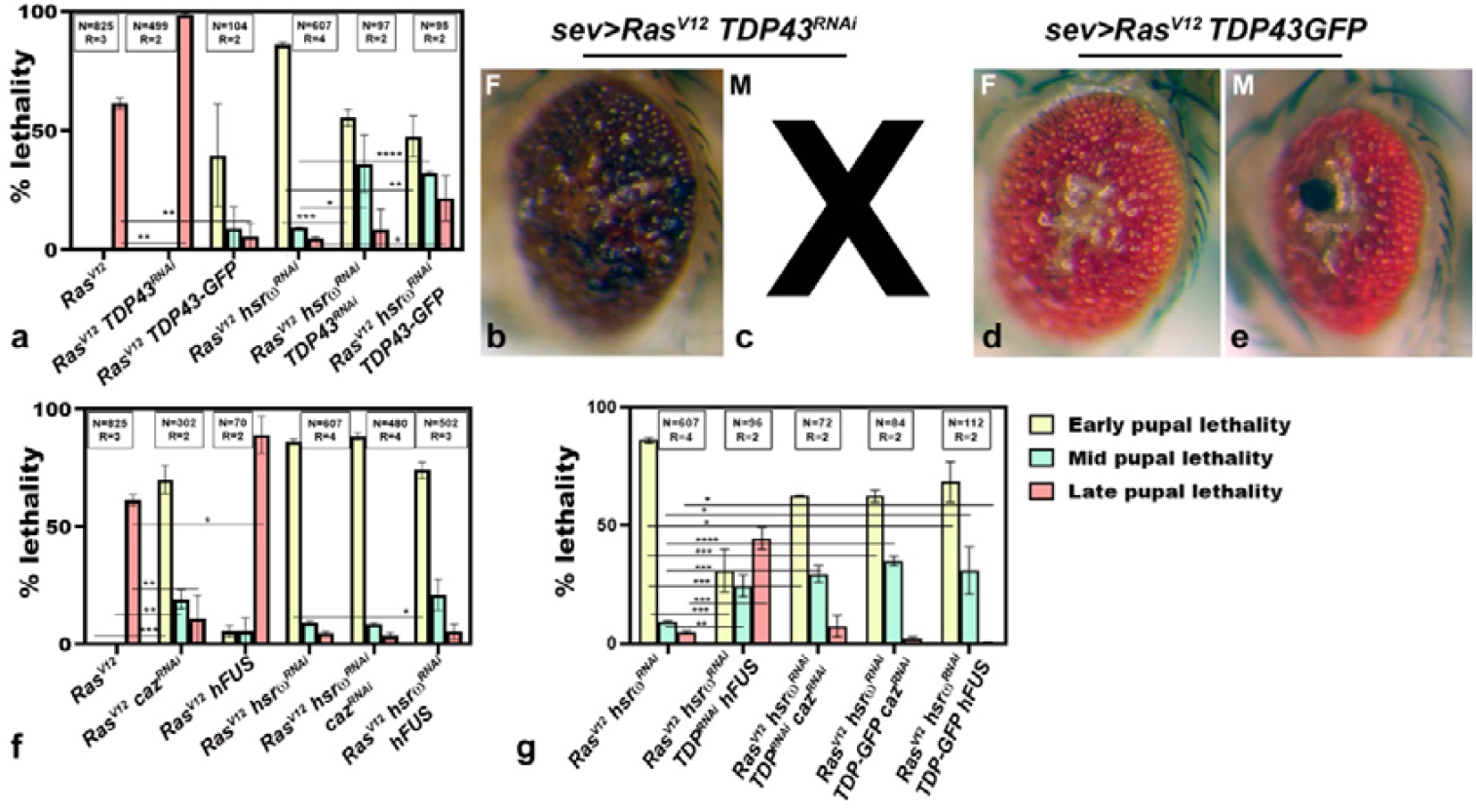
Complex modulatory effects of altered levels of TBPH and/or Caz on *sev*>*Ras*^*V12*^ expression induced eye roughening and pupal lethality. **a, f** and **g**. Histograms showing mean (± S.E) percent death (Y-axis) at early, mid or late pupal stage (color code on right of g) in different genotypes (X-axis); ‘N’ and ‘R’ values in boxes in **a, f** and **g** indicate numbers of progenies and biological replicates, respectively; horizontal lines connecting different bars indicate the pair-wise comparisons of significance of differences. With *, ** and *** indicating P<0.05, 0.01 and 0.001, respectively, on 2-tailed unpaired t-test. **b-e**. Eyes of adult female (**b, d**) and male (**c, e**) flies of genotypes indicated on top; **X** in **c** indicates complete absence of male flies of this genotype.

Down-regulation of Caz/dFus (*sev*>*Ras*^*V12*^ *caz*^*RNAi*^) or over-expression of wildtype hFUS (*sev*>*Ras*^*V12*^ *hFUS*) in *sev*>*Ras*^*V12*^ background caused complete pupal lethality with *sev*>*Ras*^*V12*^ *caz*^*RNAi*^ pupae suffering high early pupal lethality while many *sev*>*Ras*^*V12*^ *hFUS* progeny developed up to late pupal stage before dying (Fig.2f).

Changes in Caz/hFUS levels in *sev*>*Ras*^*V12*^ *hsr*ω^*RNAi*^ background (Fig. 2f) did not alter the high early pupal lethality with no emergence of adult fly seen in *sev*>*Ras*^*V12*^ *hsr*ω^*RNAi*^.

We next examined effects of co-alterations in levels of TDP-43 and Caz in *sev*>*Ras*^*V12*^ *hsr*ω^*RNAi*^ background (Fig. 2g). Down-regulation of TDP-43 in conjunction with over-expression of hFUS reduced the high early pupal death seen in *sev*>*Ras*^*V12*^ *hsr*ω^*RNAi*^ progenies, although none of the *sev*>*Ras*^*V12*^ *hsr*ω^*RNAi*^ *TDP*^*RNAi*^ *hFUS* pupae emerged. Pupal death in progeny with up-regulated TDP-43 and down-regulated Caz (*sev*>*Ras*^*V12*^ *hsr*ω^*RNAi*^ *TDP-43-GFP caz*^*RNAi*^) or with both TDP-43 and Caz down-regulated (*sev*>*Ras*^*V12*^ *hsr*ω^*RNAi*^ *TDP*^*RNAi*^ *caz*^*RNAi*^) was similar to that in *sev*>*Ras*^*V12*^ *hsr*ω^*RNAi*^ progeny. Co-over-expression of TDP-43 and hFUS in *sev*>*Ras*^*V12*^ *hsr*ω^*RNAi*^ background also did not significantly modify the high early pupal death phenotype. No adults emerged in any of these genotypes (Fig. 2g).

In summary (Table 1), the exaggeration of *sev*>*Ras*^*V12*^ expression induced damage by *hsr*ω^*RNAi*^ is generally epistatic to any modulatory effect of changes in TBPH or Caz since none of them permitted adult emergence.

**Table 1.**
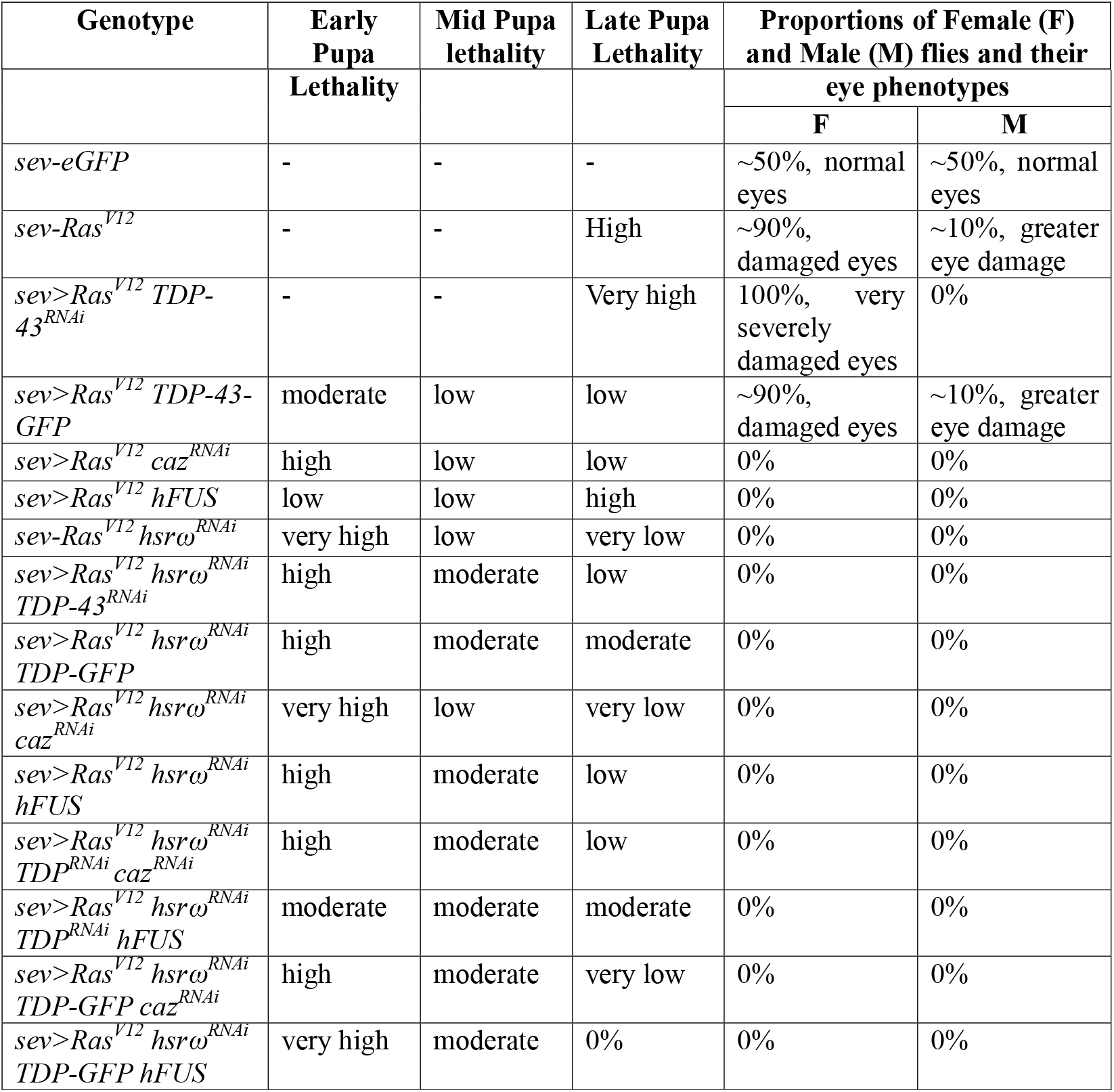
Summary of pupal lethality and sex-dependent differences in adult emergence and eye phenotypes of emerging adults (Figs. 1c, 2a, f and g)

### 3.3. Sub-cellular localizations of TDP43 and levels of Caz in eye discs modulated by *sev*>*Ras*^*V12*^ expression

Sub-cellular localization of TBPH hnRNP plays a crucial role in maintaining its regulatory functions [29, 32, 33]. Eye discs of 120hr larvae (Fig.3 a-d”) showed sex-dependent differences in nucleo-cytoplasmic distribution of TBPH in control (*sev*>*eGFP*) and *sev*>*Ras*^*V12*^ eye discs although the total TBPH immunostaining intensity did not show any genotype-or sex-dependent differences in developing ommatidia in larval eye discs (Fig.3e). As revealed by intensity profile plots (Fig. 3a’’’-d’’’) TBPH appeared to be nearly similarly distributed in nuclear and cytoplasmic areas of ommatidial cells in *sev*>*eGFP* female eye discs (Fig. a’’’) but in *sev*>*eGFP* male eye discs (Fig. b’’’), TBPH was more pronounced in cytoplasm than in nucleus. Interestingly, TBPH staining in *sev-Gal4* expressing GFP-positive cells was more pronounced in cytoplasm with proportionate decline in nuclei in *sev*>*Ras*^*V12*^ female larval eye discs (Fig. 3c’’’) while that in *sev*>*Ras*^*V12*^ male eye discs (Fig. 3d’’’) remained nearly similar to that in *sev*>*eGFP* male eye discs (Fig. 3a’’’). Quantification of TBPH in R7 nuclei (Fig. 3f) also confirmed that while the nuclear TBPH remained generally similar in male *sev*>*eGFP* and *sev*>*Ras*^*V12*^ eye discs, there was a significant reduction in nuclear TBPH in R7 cells of *sev*>*Ras*^*V12*^ female when compared with that in *sev*>*eGFP* female eye discs. It is notable that in both the genotypes, the R7 nuclear content of TBPH in female eye discs was significantly greater than in corresponding male eye discs (Fig. 3f)

**Fig. 3.**
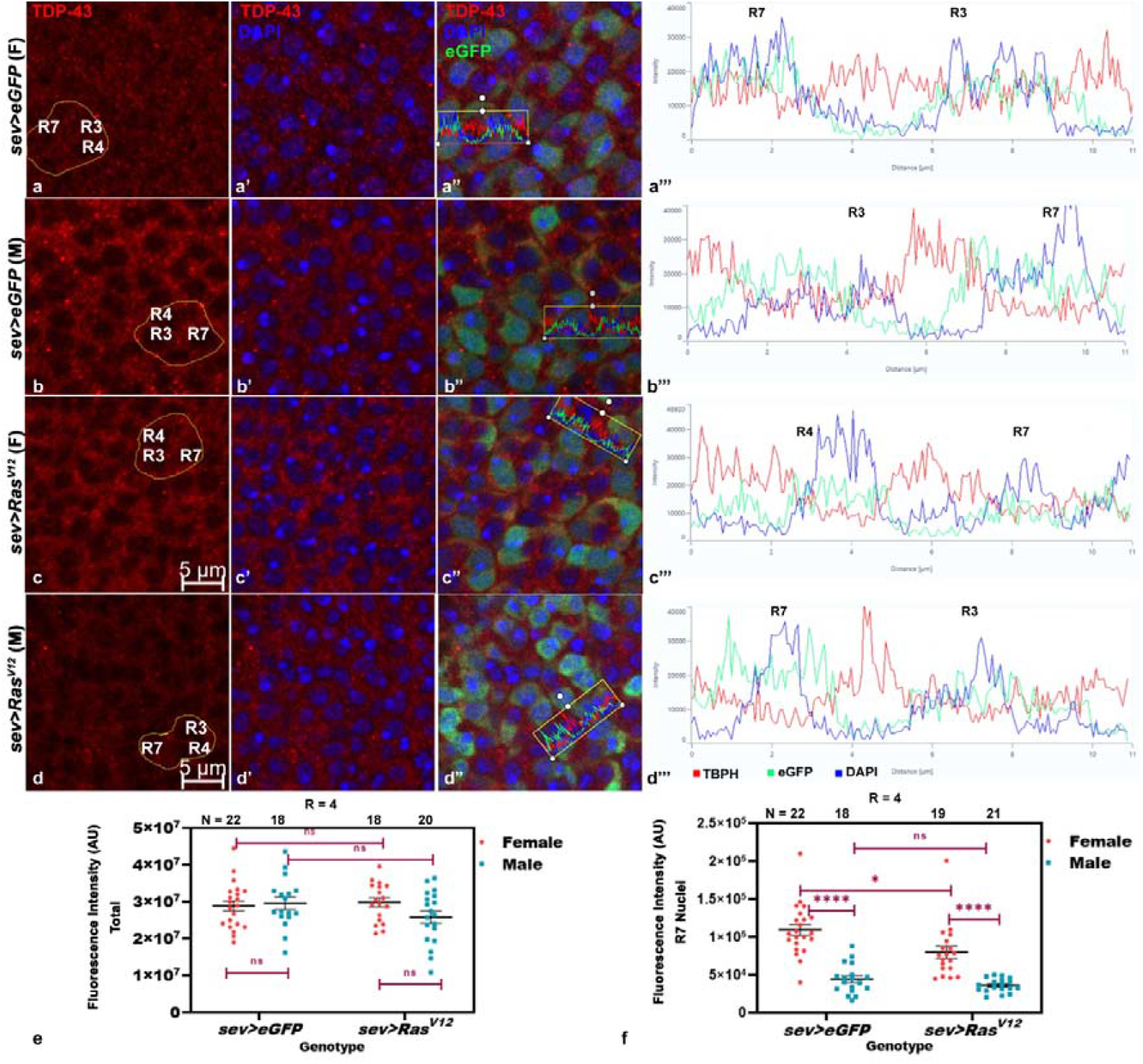
*sev*>*Ras*^*V12*^ expression alters nucleo-cytoplasmic distribution in female but not in male eye discs. **a-d”**. Confocal optical sections of proximal region of 120 hr eye discs showing localization of TBPH (red), nuclei (DAPI, blue) and eGFP (green) in male (M) and female (F) *sev*>*eGFP* (**a-b”**) and *sev*>*Ras*^*V12*^ (**c-d”**) larvae; **a’’’**-**d’’’** are intensity profile plots of TBPH (red), GFP (green) and DAPI (blue) fluorescence intensities along the scan lines shown in **a”, b”, c”** and **d”**, respectively; the R7 and R3/R4 rhabdomeres whose fluorescence intensities were scanned are marked on **a-d** as well as on the scan profiles in **a’’’-d’’’. e-f**. Dot plots of TBPH fluorescence intensities (Y-axis, arbitrary units) in defined areas of imaginal discs (**e**) or in R7 nuclei (**f**) in different genotypes (X-axis); **R** and **N** in **e** and **f** represent, respectively, biological replicates and numbers of discs examined for each genotype and gender (colour codes given on right of each plot).

Caz/dFus (*Drosophila* homolog of human FUS) hNRNP was more abundant in eye discs of male *sev*>*eGFP* larvae than in female (Fig. 4a-b’ and e). Interestingly, following *sev*>*Ras*^*V12*^ expression, levels of Caz were significantly down-regulated in both male and female eye discs (Fig. 4 c-d’); the decrease in male eye discs was proportionately greater since the difference in Caz levels in *sev*>*Ras*^*V12*^ female and male eye discs was not statistically significant, although the levels in male discs continued to appear slightly higher than in corresponding female discs (Fig. 4e).

**Fig. 4.**
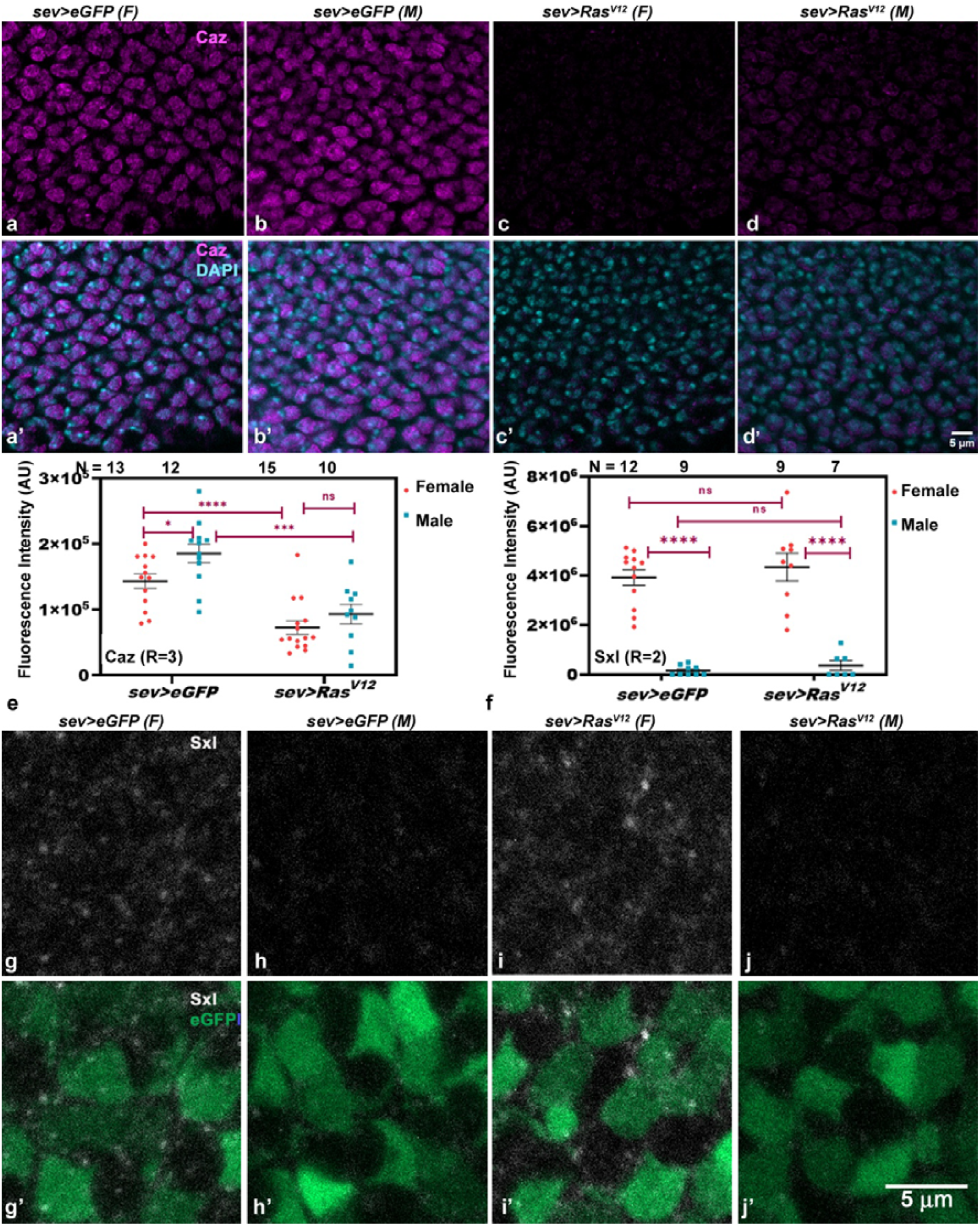
*sev*>*Ras*^*V12*^ expression inhibits Caz levels but does not affect Sxl. **a-d’**. Confocal optical sections of the proximal-posterior region of 120hr eye discs of *sev*>*eGFP* female (**a-a’**) and male (**b-b’**) and *sev*>*Ras*^*V12*^ female (**c-c’**) and male (**d-d’**) larvae immunostained for Caz (magenta in **a-d’**); DAPI is shown in cyan in **a’-d’**; scale bar in **d’** denotes 5µm and applies to **a-d’. e-f**. Dot plots of fluorescence intensities (Y-axis, arbitrary units) of Caz (**e**) and Sxl (**f**) in female and male *sev*>*eGFP* and *sev*>*Ras*^*V12*^ eye discs (X-axis); **R** and **N** (on top of each dot column) indicate the numbers of biological replicates and eye discs examined, respectively. **g-j’**. Confocal optical sections of the posterior region of 120hr eye discs of *sev*>*eGFP* female (**g-g’**) and male (**h-h’**), and *sev*>*Ras*^*V12*^ female (**i-i’**) and male (**j-j’**) larvae immunostained for Sxl (grey); eGFP is shown in green; scale bar in **j’** indicates 5µm and applies to **g-j’**

We next examined status of the key sex-determinant factor Sxl [34]. The active form of Sxl is expressed only in females and triggers female development by silencing the male specific traits [34-36]. As shown in Fig. 4f-j’, no significant difference was discernible in the total amount of Sxl (Fig. 4f) or its sub-cellular distribution between *sev*>*eGFP* and *sev*>*Ras*^*V12*^ female eye discs (Fig. 4g-g’, i-i’). As expected, Sxl was nearly undetectable in male eye discs of either genotype (Fig. 4f, h-h’ and j-j’).

### 3.4. Ras^V12^ expression affects the microtubule binding protein Futsch in eye discs

hnRNP TBPH/TDP-43 is known to regulate translation of microtubule binding protein Futsch directly by interacting with its mRNA through its RNA binding capacity [37-39]. Since the sub-cellular TBPH was affected in *sev*>*Ras*^*V12*^ expressing female eye discs (Fig. 3), we examined distribution and levels of Futsch in eye discs. The distribution and levels of Futsch were comparable in female and male *sev*>*eGFP* larval eye discs (Fig. 5a-b” and e) and were substantially reduced in female as well as male eye discs following sev>*Ras*^*V12*^ expression (Fig. 5c-d” and e).

**Fig. 5.**
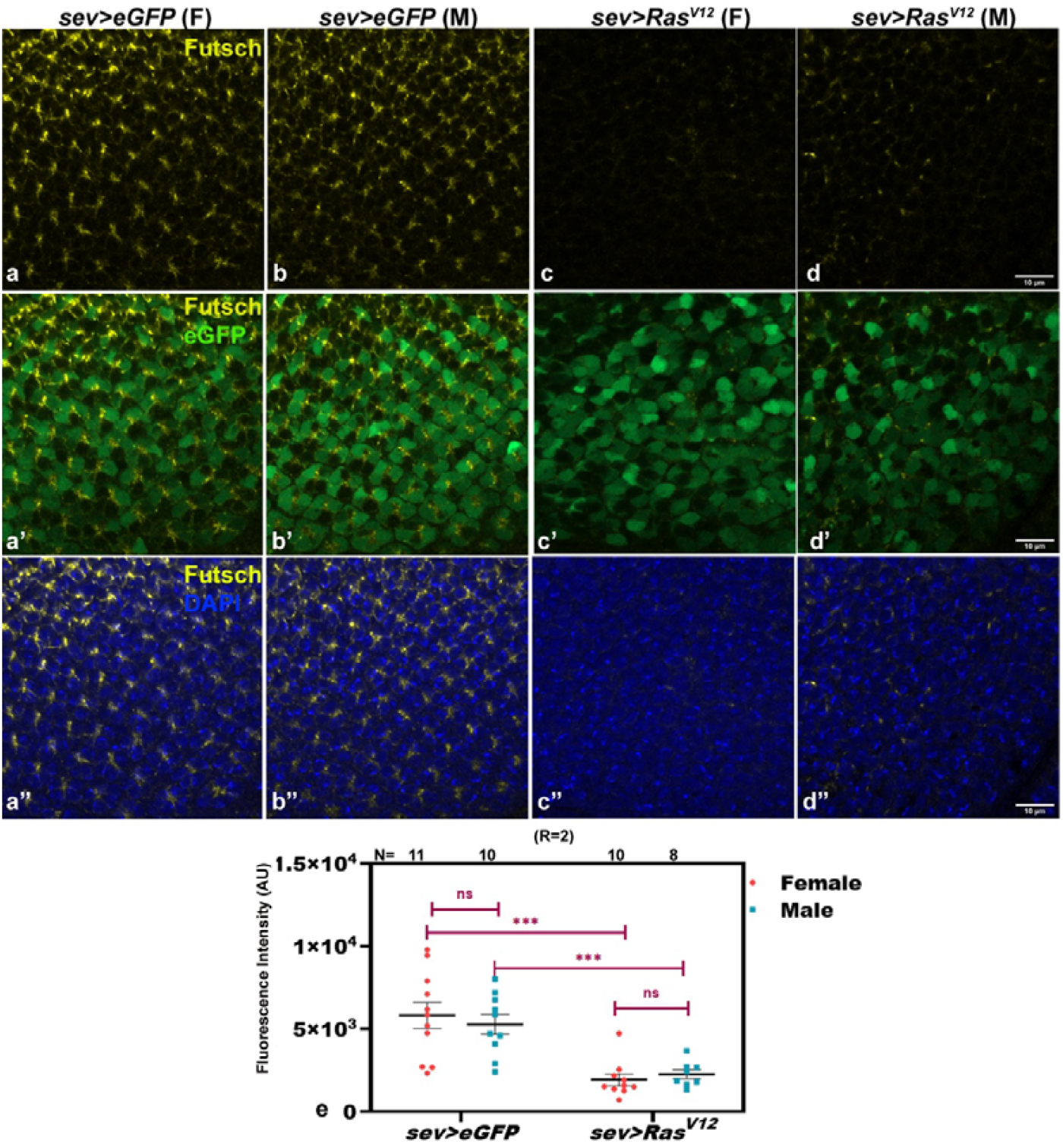
*sev*>*Ras*^*V12*^ expression substantially inhibits Futsch in eye discs. **a-d”**. Confocal optical sections showing distribution of 22C10 antibody stained Futsch in proximal posterior regions of 120hr larval eye discs in *sev*>*eGFP* female **(a-a”)** and male **(b-b”)**, *sev*>*Ras*^*V12*^ female **(c-c”)** and male **(d-d”)** larvae; eGFP and DAPI fluorescence are shown in green and blue, respectively. **e**. Dot-plots of Futsch fluorescence intensity (Y-axis) in female and male eye discs in different genotypes (X-axis)

### 3.5. Fasciclin II (Fas2), a dose dependent negative regulator of EGFR signaling, is elevated in *sev*>*Ras*^*V12*^ females but not males

FasII, a cell adhesion molecule, is also involved in EGFR signaling regulation since high and low to moderate levels of FasII regulate EGFR signaling negatively and positively, respectively, through direct interaction with EGFR receptor and affecting its phosphorylation [40-42]. FasII levels and sub-cellular distribution were comparable in *sev*>*eGFP* female and male eye discs (Fig. 6a-b” and e). Significantly, levels of FasII on cell membranes were elevated in *sev*>*Ras*^*V12*^ female eye discs, (>1.5-fold increase) especially around the GFP positive cells, but in some cases in non-GFP expressing adjacent cells as well (Fig.6 c-c” and e). However, *sev*>*Ras*^*V12*^ expression in male eye discs did not affect FasII expression (Fig.6 d-d” and e).

**Fig. 6.**
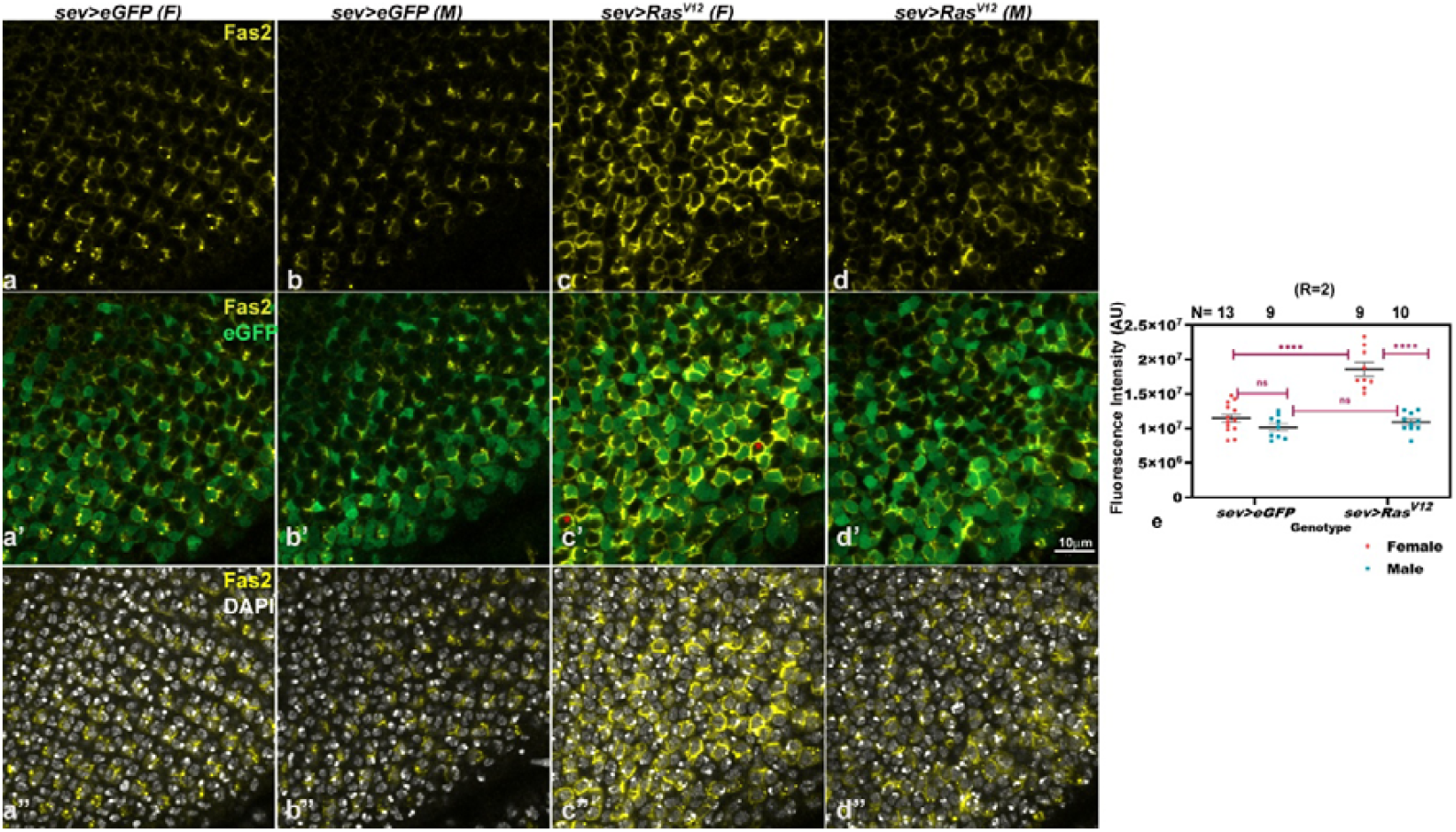
*sev*>*Ras*^*V12*^ expression elevates Fas2 levels only in female eye discs. **a-d”**. Confocal single optical sections showing proximal regions of 120 hr larval eye discs immunostained with FasII antibody (yellow) in *sev*>*eGFP* female **(a-a”)** and male **(b-b”)**, *sev*>*Ras*^*V12*^ female **(c-c”)** and male **(d-d”)**; red asterisks (**\*)** in **c’** mark non-eGFP (green) expressing cells with elevated FasII levels; DAPI fluorescence is in gray. Scale bar in **d’** denotes 10µm and applied to all the images. **e**. Dot plots of Fas2 fluorescence intensity (Y-axis) in female and male eye discs of different genotypes (X-axis); **R** and **N** indicate, respectively, the biological replicates and numbers of eye discs examined.

### 3.6. Greater increase in cell numbers following *sev*>*Ras*^*V12*^ expression in male than in female eye discs

In the earlier study, [1] reported that *sev*>*Ras*^*V12*^ expression was accompanied by additional R7 cells in ommatidial units. In the present study, we used Gal4 Technique for Real-time and Clonal Expression or G-TRACE [43] analysis to examine if the Gal4 expressing cells and their clonal/lineage cells follow different dynamics in male and female eye discs. In this method, the *sev-Gal4* expressing cells show red fluorescence (RFP) while their lineage cells, which may no longer express the *sev-Gal4* transgene, exhibit green fluorescence (GFP) but the lineage cells continuing to express *sev-Gal4*, show red as well as green fluorescence. Our results revealed that the numbers of RFP-expressing cells in *sev*>*G-TRACE* and *sev*>*G-TRACE Ras*^*V12*^ female eye discs remained similar (Fig. 7a, c e) although the GFP-expressing *sev-Gal4* lineage cells were more abundant in *sev*>*G-TRACE Ras*^*V12*^ female eye discs (Fig. 7a’, c’, e). In contrast, the RFP as well as GFP-expressing cells were significantly more in *sev*>*G-TRACE Ras*^*V12*^ than in *sev*>*G-TRACE* male eye discs (Fig. 7b, b’, d, d’, e). Apparently, the greater increase in Ras level in *sev*>*Ras*^*V12*^ male eye discs (Fig.1) triggers greater number of *sev*>*Ras*^*V12*^ lineage cells, which results in more severe eye roughening in *sev*>*Ras*^*V12*^ males (Fig.1).

**Fig. 7.**
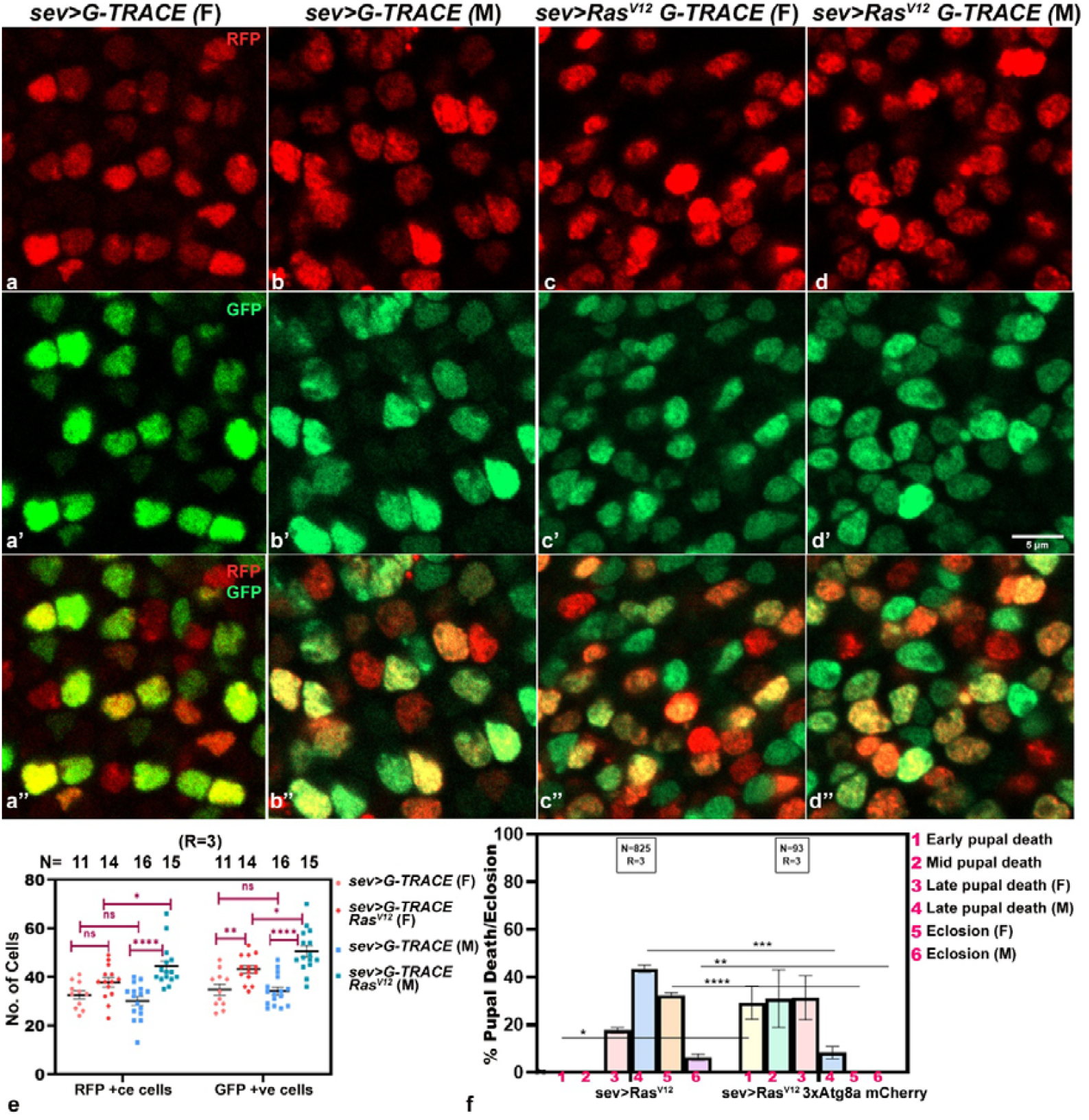
Eye discs expressing *sev-Gal4* driven *Ras*^*V12*^ have more cells in male larvae. **a-d”**. Confocal single optical sections showing proximal region of 120 hr larval eye discs of *sev*>*G-TRACE* female **(a** and **a’’)**, *sev*>*G-TRACE* male **(b** and **b’’)**, *sev*>*G-TRACE Ras*^*V12*^ female **(c** and **c’’)** and *sev*>*G-TRACE Ras*^*V12*^ male **(d** and **d’’)** eye discs with RFP **(a-d)** and GFP positive cells **(a’-d’)**; **a’’-d’’** show combined RFP and GFP fluorescence signals; scale bar in **d’** represents 5µm and applies to all the images. **e**. Dot plots of mean numbers (± S. E., Y-axis) of RFP and GFP positive cells in *sev*>*G-TRACE* and *sev*>*G-TRACE Ras*^*V12*^ female and male eye discs (X-axis; colour code indicated on right of the image); **R** and **N** on top indicate the numbers of independent replicates and eyes discs examined in each sample. **f**. Histogram showing mean (± S.E.) precent pupal lethality and eclosion (Y-axis) of *sev*>*Ras*^*V12*^ and *sev*> *Ras*^*V12*^ *3x Atg8a-mcherry* progenies (X-axis); **R** and **N** in boxes on top indicate numbers of progenies and biological replicates, respectively.

### 3.7. The s*ev*>*Ras*^*V12*^ expression induced pupal lethality is enhanced with overexpression of Atg8a

Autophagy is known to be required for tissue overgrowth in Ras^V12^ background [44]. Since the earlier [1] and our above data show that *sev*>*Ras*^*V12*^ eye discs carry enhanced cell numbers, especially in males (Fig. 7a-e), we examined if enhanced autophagy following expression of *3x Atg8a mCherry* transgene (with its own endogenous promoter) in *sev*>*Ras*^*V12*^ eye discs would exaggerate the pupal lethality, more so in males than in females. Indeed, overexpression of Atg8a in *sev*>*Ras*^*V12*^ background, enhanced early and mid-stage pupal death (Fig.7f). Since among those dying as late pupae there were far fewer males than females, the dead early and mid-pupae would have included more males than females. The enhanced pupal lethality following Atg8a over-expression in *sev*>*Ras*^*V12*^ background is also reflected in complete absence of emergence of male as well as female adults (Fig.7f). It is noted that the *w*^*1118*^; *+/+; 3x Atg8a-mcherry* stock by itself does not show any pupal lethality or other phenotypic effect due over-expression of Atg8a under its endogenous promoter.

## 4. Discussion

Following earlier observations from our laboratory [1, 2] that the systemic phenotypes resulting from over-expression of activated Ras in *sevenless* expressing cells in *Drosophila* larval eye discs (*sev*>*Ras*^*V12*^) and their exaggeration by co-perturbations in levels of the *hsr*ω lncRNAs, we now show these effects to be sex-biased with males being more sensitive. It may be noted that the earlier [45] report that the *sev-Gal4* driver expresses also in 8 pairs of dorso-medial neurons in the ventral ganglia besides the specific cells in developing eye discs was erroneous since our subsequent study [46] revealed that the earlier claimed expression of *sevGal4* in the ventral ganglia neurons was actually due to the default expression of the *UAS-GFP* reporter transgene used earlier. Consequentially, the adverse local and systemic phenotypes in *sev*>*Ras*^*V12*^ individuals [1, 2] seem to originate primarily because of Ras^V12^ expression within the *sevenless* expressing eye disc cells.

Earlier study [1] indicated that several hnRNP genes were commonly affected in *sev*>*Ras*^*V12*^ and *sev*>*Ras*^*V12*^ *hsr*ω^*RNAi*^ eye discs. Our present genetic interaction studies (Table 1) indeed showed that up-as well as down-regulation of TBPH and /or Caz, two key hnRNPs involved in multiple regulatory processes in cells [11, 47-54], varyingly modulated the sex-biased rough eye phenotype and pupal lethality associated with *sev*>*Ras*^*V12*^ and *sev*>*Ras*^*V12*^ *hsr*ω^*RNAi*^ genotypes.

Ras signaling is a key pathway regulating multiple biological processes ranging from cell proliferation to cell death [55]. The significantly greater increase in *sev*>*Ras*^*V12*^ lineage cells in male eye discs would disrupt the development to a greater extent than in female and therefore, result in the observed male-biased greater eye roughness and pupal lethality. An earlier study [44] reported that directed reduction of autophagy reduces Ras^V12^ mediated tissue overgrowth. Our finding that enhanced autophagy following over-expression of Atg8a in *sev*>*Ras*^*V12*^ background further aggravated the male pupal lethality also indicates that the male-biased exaggerated phenotypes are related to the greater Ras levels in *sev*>*Ras*^*V12*^ males than in sib-females.

As summarized in Table 2, we found differences in levels and/or sub-cellular distributions of Caz and TBPH between *sev*>*eGFP* (control) male and female larval eye discs. TBPH was more cytoplasmic than nuclear and Caz more abundant in male *sev*>*eGFP* (control) than in female larval eye discs. Since functional dynamics of the diverse hnRNPs in *Drosophila* cells are majorly interconnected through the omega speckles organized by nuclear *hsr*ω lncRNAs [26, 27, 56, 57], we believe that the network and regulatory feedback effects of these differences and the well-known absence of Sxl in males [58] together culminate in the differential levels of Ras in *sev*>*Ras*^*V12*^ male and female larval eye discs.

**Table 2.** Summary of changes in protein levels and their sub-cellular localization following *sev*>*Ras*^*V12*^ expression.

| Protein | Protein levels |  |  |  | Sub-cellular localization |  |  |  |
| --- | --- | --- | --- | --- | --- | --- | --- | --- |
|  | *sev>eGFP* |  | *sev>Ras<sup>V12</sup>* |  | *sev>eGFP* |  | *sev>Ras<sup>V12</sup>* |  |
|  | F | M | F | M | F | M | F | M |
| Ras | + | + | ++ | +++ | Cellular periphery with occasional puncta | Cellular periphery with occasional puncta | Elevated around GFP+ve and adjacent GFP-ve cells | **Greater elevation** around GFP+ve and adjacent GFP-ve cells |
| TBPH | + | + | + | + | Distributed in nucleus and cytoplasm | **More cytoplasmic** | **More in cytoplasm and less in nucleus** | As in *sev>eGFP* male |
| Caz | + | ++ | - | - | Nucleus | Nucleus | Nuclear but reduced | Nuclear but reduced to nearly the level in *sev>Ras<sup>V12</sup>* female (**greater reduction**) |
| Sxl | + | - | + | - | Diffuse in nucleus and cytoplasm with a few puncta | Absent | Diffuse in nucleus and cytoplasm with a few puncta | Absent |
| Futsch | + | + | - | - | Cell periphery | Cell periphery | Significantly reduced | Significantly reduced |
| FasII | + | + | ++ | + | Cell periphery; more prominent on *sev* expressing cells | Cell periphery; more prominent on *sev* expressing cells | **Significant elevation around *sev* expressing cells** | As in *sev>eGFP* male |

Significant differences between female and male eye discs of comparable larval genotypes marked in bold.

TBPH/TDP-43 and Caz/Fus are very versatile regulators of RNA processing, transport, and translation, and major players in various neurodegenerative disorders because of their phase-separation dependent aggregation following interactions with each other, diverse RNAs and other hnRNPs [48-54]. Elevation in activated Ras and the consequent elevation in MAPK activity hyper-phosphorylates TBPH, which reduces its binding with RNA and leads to TBPH moving from nucleus to cytoplasm [23, 24, 59] as observed in *sev*>*Ras*^*V12*^ female larval eye discs. The greater cytoplasmic TBPH seen in male *sev*>*eGFP* (control) eye discs appears to be developmentally regulated and associated with absence of Sxl and higher Caz. This default situation may not lead to further increase in cytoplasmic TBPH even when Ras levels increase in *sev*>*Ras*^*V12*^ male larval eye discs.

Caz is reported to negatively regulate EGFR signaling [25]. Our results suggest that EGFR/JNK signaling may also negatively regulate Caz since elevation in Ras levels in eye disc cells was accompanied by substantial reduction in Caz. In agreement, co-expression of *caz*^*RNAi*^ in *sev*>*Ras*^*V12*^ resulted in substantially enhanced early pupal death with no eclosion while over-expression of *hFUS* delayed the pupal death to later stages, although no adults could emerge. Presumably, excess hFUS contributed its own toxicity so that compared to the limited eclosion of *sev*>*Ras*^*V12*^, none of the *sev*>*Ras*^*V12*^ *hFUS* pupae eclosed.

Interactions between TBPH, Caz, and with Sxl in female cells maintain their correct sub-cellular localizations and functions [31, 34-36, 60]. Since down-regulation of Caz or its mutation is often associated with cytoplasmic migration of nuclear TBPH [30, 31, 60], the significantly reduced Caz levels in *sev*>*Ras*^*V12*^ eye discs likely contributes to the reduced nuclear and greater cytoplasmic TBPH in *sev*>*Ras*^*V12*^ eye discs; this effect was noticeable only in *sev*>*Ras*^*V12*^ female eye discs since in male *sev*>*eGFP* eye discs, TBPH was already less abundant in nuclei.

The significantly higher levels of Fas2 in female eye discs expressing *sev*>*Ras*^*V12*^ is significant since Fas2 and EGFR/Ras signaling are involved in auto-regulatory loop in which Fas2 is a cell autonomous EGFR activator at low and moderate expression levels but an EGFR repressor at high levels of expression [40-42, 61, 62]. The observed elevation in Fas2 levels only in *sev*>*Ras*^*V12*^ female eye discs appears to be related to interactions between Sxl, TBPH and other hnRNPs in these cells. Mef2 transcription factor, a negative regulator of FasII levels [63], is regulated by hnRNPs like Sxl, hnRNP L/Smooth, HnRNP D/AUF1, TBPH, Caz, Hrb87F [36, 64, 65], Nej/CBP300, MAPK etc [66, 67]. The hnRNP L regulates transcription and splicing patterns of Mef2 while hnRNP D promotes Mef2 translation [64]. Since activities of TBPH in RNA splicing, transport etc depend upon its nuclear localization and its RNA binding affinity, its movement from nucleus to cytoplasm and the reduction in Caz in *sev*>*Ras*^*V12*^ female eye discs may disrupt the Mef2 activity. Additionally, binding of CBP/P300 transcription regulator with Mef2 is reported to repress Mef2 activity, which in turn stimulates Pumilio, a negative regulator of EGFR signaling [68, 69]. CBP/P300 physically interacts with several hnRNPs [70]; this makes it possible that the alterations of some of the key hnRNPs in *sev*>*Ras*^*V12*^ female eye discs and the presence of Sxl therein further disrupts Mef2 activity. Together this results in the observed differences in Fas2 levels in the two sexes. High Fas2 would inhibit EGFR and thus reduce Ras levels in *sev*>*Ras*^*V12*^ female eye discs. A similar autoregulation of Ras and Fas2 does not occur in *sev*>*Ras*^*V12*^ male eye discs since TBPH is located in cytoplasm by default, which does not change following *sevGal4* driven Ras^V12^ expression.

We speculate that Sxl, which is present only in female cells, plays a key role in TBPH-CBP-Mef2 regulatory events since Sxl interacts with diverse hnRNPs either directly or through the *hsr*ω transcripts [36, 57, 71]. It is known that interactions between Hrp48/Hrb27C and Sxl regulate sex-specific Notch signaling during eye development in *Drosophila* [72]. In a similar manner, interactions of Sxl may determine the regulatory effects of various hnRNPs and transcription factors in modulating Fas2 levels and Ras signaling. Such differential interactions of hnRNPs and their targets in the two sexes finally cause greater levels of Ras in *sev*>*Ras*^*V12*^ expressing male eye discs and the consequent sex-biased deleterious consequences.

The Futsch protein of *Drosophila* is a microtubule associated protein that is essential for organization of dendritic, axonal, and nerve-terminal cytoskeleton [73]. Its drastic reduction in *sev*>*Ras*^*V12*^ expressing eye discs was associated with disruption in organization of axons in the optic nerve as well as the systemic effect is also seen in axons of the ventral ganglia in the central nervous system (not shown). The significant down-regulation of Futsch in *sev*>*Ras*^*V12*^ expressing eye discs appears to be due to the altered localization and consequent reduced RNA binding affinity of TBPH since it is known to physically interact with Futsch mRNA and regulate its translation [37-39]. Although Futsch is not known to affect EGFR/Ras signaling, the disruption in photoreceptor axon organization is expected to indirectly affect the EGFR signaling required for rhabdomere development.

It is interesting that the sex-dependent sensitivity is related to sex-determination pathway and/or sex-chromosome-linked genetic factors in most cases [3-9]. In the present case also, the gender-associated differences in interactions of hnRNPs and the main sex-determination protein Sxl appear to be the causal factors for the male-biased deleterious consequences of over-expression of activated Ras in a subset of developing eye cells. The regulatory networks of Ras-signaling as well as the hnRNPs are extensive, multi-component and developmental stage-, cell- and tissue-specific interactions. Present results illustrate how apparently small perturbations in levels and activities of a component of these network can get biologically magnified and result in severe local as well as systemic effects.

## Acknowledgement

We thank Bloomington Drosophila Stock Center; USA, Prof. Girish Ratnaparkhi (IISER Pune), Dr. Udaybhan Pandey (Louisiana State University), Dr. Moushami Mallik (Germany), Dr. Bhupendra Shravage; Agarkar Research Institute (Pune), and Dr. Daniella Zarnescu (University of Arizona) for fly stocks and antibodies. We would also like to express sincere appreciation to Dr. Bama Charn Mondal (Dept Zoology), ISLS, and CDC of BHU for instrumentation facility.

## Author contributions

Vanshika Kaushik: Conceptualization, Data curation, Formal analysis, Investigation, Methodology, Validation, Visualization, and Writing original draft and review and editing.

S.C. Lakhotia: Conceptualization, Project administration, Resources, Supervision, Validation, Writing and review-editing.

## Data availability

All relevant data can be found within the text.

